# Honey bee parasitic mite contains the sensory organ expressing ionotropic receptors with conserved functions

**DOI:** 10.1101/505529

**Authors:** Jing Lei, Qiushi Liu, Tatsuhiko Kadowaki

## Abstract

Honey bee parasitic mites (*Tropilaelaps mercedesae* and *Varroa destructor*) detect temperature, humidity, and odor but the underlying sensory mechanisms are poorly understood. To uncover how *T. mercedesae* responds to environmental stimuli inside a hive, we identified the sensilla-rich sensory organ on the foreleg tarsus. The organ contained four types of sensilla, which may respond to different stimuli based on their morphology. We found the forelegs were enriched with mRNAs encoding sensory proteins such as ionotropic receptors (IRs) and gustatory receptors (GRs), as well as proteins involved in ciliary transport. We also found that *T. mercedesae* and *Drosophila melanogaster* IR25a and IR93a are functionally equivalent. These results demonstrate that the structures and physiological functions of ancient IRs have been conserved during arthropod evolution. Our study provides insight into the sensory mechanisms of honey bee parasitic mites, as well as potential targets for methods to control the most serious honey bee pest.

## Introduction

The number of managed honey bee colonies has declined across North America and Europe in recent years ^1^. Pollination by honey bees is critical for maintaining ecosystems and producing many agricultural crops ^2,3^ Prevention of honey bee losses has, therefore, become a major issue in apiculture and agriculture. Although there are many potential causes for the observed declines, ectoparasitic mites are considered to be major threats to the health of honey bees and their colonies ^1,4^ *Varroa destructor* is present globally (except Australia) and causes both abnormal brood development and brood death in honey bee colonies ^5^. The mites feed on hemolymph and also spread honey bee viruses, particularly deformed wing virus (DWV) ^6,7^ In many Asian countries, another honey bee ectoparasitic mite, *Tropilaelaps mercedesae*, is also prevalent in *Apis mellifera* colonies ^8,9^ These two emerging parasites of *A. mellifera* share many characteristics ^10^. For example, they have similar reproductive strategies ^11^ and both are vectors for DWV ^12-15^. As a result, *T. mercedesae* or *V. destructor* infestations have similar negative impacts on *A. mellifera* colonies ^16–18^. Although *T. mercedesae* is currently restricted to Asia, it has the potential to spread and establish worldwide due to the global trade in honey bees.

*V. destructor* prefers temperatures of 32 ± 2.9 °C, reproduces best at 32.5-33.4 °C, and has been shown to discriminate temperature differences of 1 °C ^19–21^. Furthermore, its reproduction also depends on humidity of 55-70 % ^22^ These results demonstrate thermo- and hygrosensation of *V. destructor* play important roles to adapt to the honey bee hive environment; nevertheless, chemoreception must be most important in the various interactions between mites and their honey bee hosts. For example, *V. destructor* prefers to parasitize nurse bees rather than foragers during its phoretic phase ^23,24^. For its reproductive stage, it locates fifth instar honey bee larva and enters the brood cell prior to capping ^25^. These behaviors are considered to be mediated by chemical cues derived from the adult bee, larva, and larval food. Since *T. mercedesae* has a very similar life cycle to *V. destructor*, both honey bee mites should be equipped with thermo-, hygro-, and chemosensation, as observed in other mite/tick (Acari) species. Accordingly, *V. destructor* was found to have a sensilla-rich sensory organ on the foreleg tarsus ^26,27^, which corresponds to Haller’s organ in ticks ^28^. Proteomic and transcriptomic characterization were conducted for the forelegs of *V. destructor*, which identified potential semiochemical carriers and sensory proteins ^29^

Ionotropic receptors (IRs) represent a subfamily of ionotropic glutamate receptors (iGluRs), which are conserved ligand-gated ion channels. IRs have specifically evolved in protostomes ^30^ and are best characterized in the fruit fly, *Drosophila melanogaster*. Most IRs are expressed in sensory neurons and function as chemoreceptors to detect various odorants and tastants ^31,32^ Recent studies have also demonstrated that IR21a, IR40a, IR68a, IR93a, and IR25a are critical for thermo- and hygrosensation, suggesting that IRs have diverse physiological roles as well as gating mechanisms ^33–36^. IR25a has the same protein domains as iGluRs, is expressed broadly in various sensory neurons, and is deeply conserved in protostomes. These findings suggest that IR25a is likely to function as a co-receptor with other IRs, similar to the OR83b pairing with other olfactory receptors (ORs).

In this study, we aimed to identify and characterize a sensilla-rich sensory organ in *T. mercedesae* using scanning electron microscopy (SEM). By comparing the transcriptomes of forelegs and hindlegs (the 2nd-4th legs), we identified potential genes that may be highly expressed in the sensory organ. Identification of this major sensory organ and its associated proteins in *T. mercedesae* inform our understanding of the mechanisms of sensory perception in honey bee parasitic mites.

## Results

### Identification of a sensilla-rich sensory organ on the foreleg tarsus of *T. mercedesae*

We observed the forelegs and hindlegs of *T. mercedesae* using SEM and found that only the foreleg tarsus contained a putative sensory organ on the dorsal side, with more than 20 sensilla of various shapes and sizes (Fig. 1A-D). Most of the sensilla were equipped with well-defined sockets (Fig. 1A). We characterized the shape of each sensillum at high magnification and found that they could be classified into four different types based on the shape: type 1 had a rough surface, e.g., #3 (Fig. 1E), type 2 had a terminal pore, e.g., #18 (Fig. 1F), type 3 included sensilla with a smooth surface, e.g., #8 (Fig. 1G), and type 4 had surface pores at various densities—sensilla #2, #7, and #10 had pores at high, medium, and low density, respectively (Fig. 1H-J). Several long sensilla were found on all legs and these are likely to be mechanosensory bristles.

**Figure 1.**
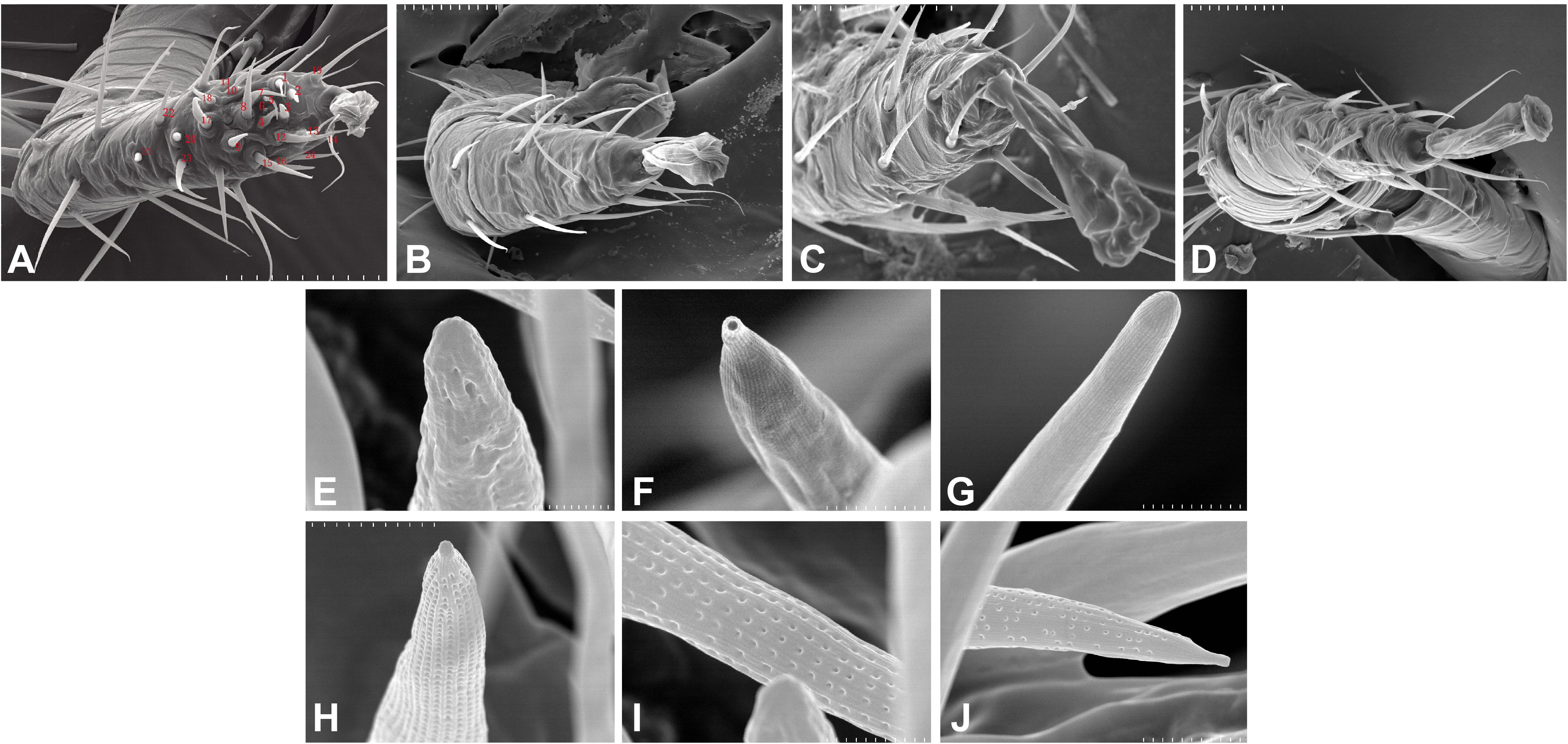
Scanning electron micrographs of *Tropilaelaps mercedesae* sensory organ. (A) The foreleg with the numbered sensilla. (B) The second leg. (C) The third leg. (D) The fourth leg. (E) Sensillum #3 with a rough surface. (F) Sensillum #18 with a terminal pore. (G) Sensillum #8 with a smooth surface. (H) Sensillum #2 with surface pores of high density. (I) Sensillum #7 with surface pores of medium density. (J) Sensillum #10 with surface pores of low density. A scale represents 50 μm in the panels A-D, 2 μm in the panels H and J, and 1 μm in the panels E-G and I.

### Identification of potential mRNAs enriched with the sensory organ

To identify potential mRNAs highly expressed in the sensory organ, we obtained RNA-seq reads from the forelegs, hindlegs, and main bodies (without legs) and then identified the differentially expressed genes (DEGs) between the forelegs and hindlegs. Since only foreleg tarsi were equipped with the sensory organs, we expected the DEGs to represent the sensory organ-associated mRNAs. We found that 46.1-83.9% of the sequence reads were aligned with the *T. mercedesae* genome (Table S1) and we used these to identify DEGs between the forelegs and hindlegs, the forelegs and main bodies, and the hindlegs and main bodies. The lists of DEGs are shown in Tables S2, S3, and S4. Tables 1, S5, and S6 indicate gene ontology (GO) terms enriched for the genes highly expressed in the forelegs compared to the hindlegs and main bodies, and the ones highly expressed in the hindlegs relative to the main bodies, respectively. For the genes highly expressed in the forelegs, many of the GO terms were associated with ion channel activity, particularly iGluR activity, as well as microtubule motor activity in the “Molecular function” category. In the “Biological process” category, GO terms related to cilium assembly, microtubule-based processes, and detection of chemical stimulus involved in sensory perception were most prevalent. All GO terms in the “Cellular component” category were related to cilium, intraciliary transport particle, and BBSome (Table 1). Several GO terms related to mitochondrial activity were also enriched in the forelegs, compared with the main bodies, and this was similar for the genes highly expressed in the hindlegs relative to the main bodies (Tables S5 and S6). These results are consistent with the finding that the sensory organ on the foreleg tarsus had many sensilla (Fig. 1) and with the ciliated sensory neurons and the expression of abundant iGluR mRNAs. It is likely that higher expression of mRNA of genes involved in mitochondrial activity in the legs relative to the main bodies would be necessary to supply energy for leg movement.

**Table 1.** GO terms enriched with genes highly expressed in the forelegs compared with hindlegs of *T. mercedesae*.

| GO ID | GO name | GO category | FDR | P-value |
| --- | --- | --- | --- | --- |
| GO:0005272 | sodium channel activity | Molecular function | 1.83E-09 | 8.77E-13 |
| GO:0004970 | ionotropic glutamate receptor activity | Molecular function | 5.02E-06 | 1.05E-08 |
| GO:0005234 | extracellularly glutamate-gated ion channel activity | Molecular function | 0.00237124 | 1.68E-05 |
| GO:0008527 | taste receptor activity | Molecular function | 0.00237124 | 1.90E-05 |
| GO:0008017 | microtubule binding | Molecular function | 0.02302381 | 2.36E-04 |
| GO:1990939 | ATP-dependent microtubule motor activity | Molecular function | 0.02706201 | 2.86E-04 |
| GO:0006814 | sodium ion transport | Biological process | 3.93E-07 | 3.78E-10 |
| GO:0050912 | detection of chemical stimulus involved in sensory perception of taste | Biological process | 0.00237124 | 1.90E-05 |
| GO:1905515 | non-motile cilium assembly | Biological process | 0.00237124 | 1.90E-05 |
| GO:0042073 | intraciliary transport | Biological process | 0.00237124 | 1.90E-05 |
| GO:0010378 | temperature compensation of the circadian clock | Biological process | 0.02706201 | 2.86E-04 |
| GO:0030990 | intraciliary transport particle | Cellular component | 0.00237124 | 1.90E-05 |
| GO:0034464 | BBSome | Cellular component | 0.01633013 | 1.60E-04 |

In addition to iGluRs, the forelegs expressed high levels of transient receptor potential channel A1 (as previously reported by Dong et al. ^37^, anoctamin-7 (TMEM16 family), and two gustatory receptors (GRs): Tm03548 and Tm05586 ^37^ (Table S2 and Fig. 2). Orthologs of these GRs were also present in *Ixodes scapularis*, but were not found in *D. melanogaster*, indicating that they are specifically expanded in the Acari lineage. Thus, the *T. mercedesae* sensory organ appears to be equipped with various sensory proteins with ion channel activity.

**Figure 2.**
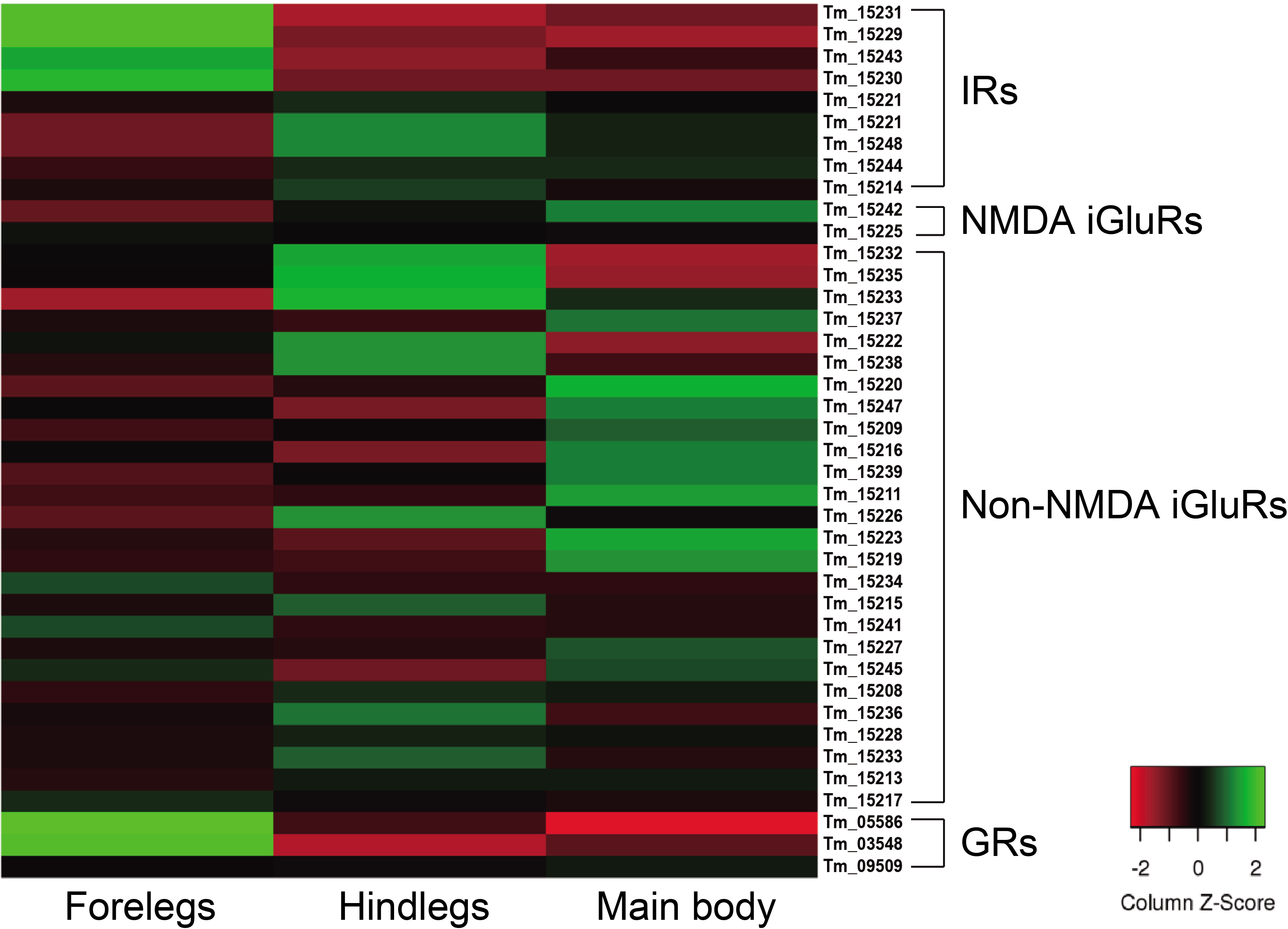
Expression of ionotropic receptors (IRs), NMDA iGluRs, non-NMDA iGluRs, and gustatory receptors (GRs) mRNAs in the forelegs, hindlegs, and main body of *T. mercedesae*. The level of expression of each mRNA in the forelegs, hindlegs, and main body is shown by a graded color (red to green) based on the counts per million mapped reads (CPM).

### Conserved sensory functions between *D. melanogaster* and *T. mercedesae* IR25a and IR93a

We previously annotated eight IR and 33 iGluR genes in the *T. mercedesae* genome and showed that two IR mRNAs, *Tm15229* and *Tm15231*, are abundantly expressed in the forelegs, using qRT-PCR ^37^. These two genes are included in the above DEGs and we also found that mRNAs for two non-NMDA iGluRs (Tm15234 and Tm15241), as well as two other IRs (Tm15230 and Tm15243), were also highly expressed in the forelegs (Fig. 2). Thus, a small fraction of iGluRs and half of IRs appear to play roles in mite sensory perception.

Based on the phylogenetic tree of *T. mercedesae* IR and iGluR genes, together with those of *D. melanogaster* and *Ixodes scapularis* ^37^, we found only two (out of eight) IRs (*Tm15229* and *Tm15231*) were conserved, having the *D. melanogaster* orthologs, *IR93a* and *IR25a*, respectively. DmIR93a and DmIR25a have been shown to play roles in temperature and humidity preferences ^33,35,36^. To test whether the sensory functions of IR93a and IR25a are deeply conserved between fruit flies and mites, we first obtained the full length cDNAs of *Tm15229* and *Tm15231* by determining both the 5’ and 3’ end sequences using RACE methods. Tm15229 (TmIR93a) and Tm15231 (TmIR25a) share the same protein domains with DmIR93a and DmIR25a, respectively (Fig. 3). Tm/DmIR25a contains the N-terminal leucine/isoleucine/valine-binding protein (LIVBP)-like domain and PBP2_iGluR domain. Meanwhile, Tm/DmIR93a contained only the PBP2_iGluR domain. The protein expression was confirmed by ectopic expression in HEK293 cells, followed by western blot (Fig. S1). We then compared the thermotactic behavior of *D. melanogaster IR93a* and *IR93a* mutants expressing *TmIR93a* under *DmIR25a-Gal4* with the wild type. Expression of *DmIR93a* and *DmIR25a* overlapped in the antennae ^35^. We also analyzed *D. melanogaster IR25a* and *IR25a* mutants expressing *DmIR25a* or *TmIR25a* under *DmIR25a-Gal4*. From our assay to test thermotactic behavior, the fraction of animals in the area with temperatures <24 °C significantly increased in both *IR93a* and *IR25a* mutants compared with the wild type; however, expression of *TmIR93a, TmIR25a*, or *DmIR25a* rescued this behavioral defect (Fig. 4A).

**Figure 3.**
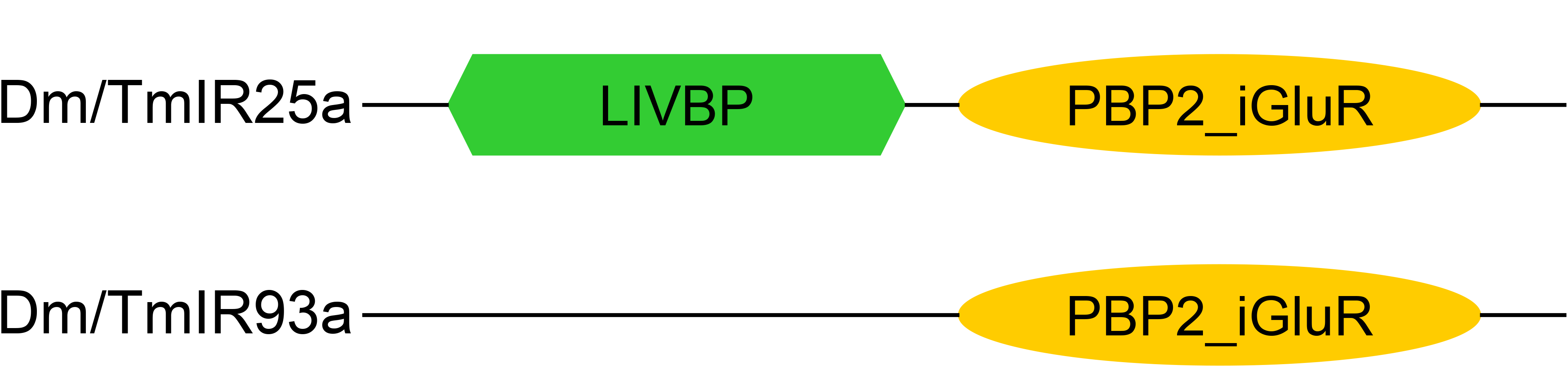
Protein domains in Dm/TmIR25a and Dm/TmIR93a. Tm/DmIR25a contains a leucine/isoleucine/valine-binding protein (LIVBP)-like domain at the N-terminus and PBP2_iGluR domain (ligand-binding and ion channel domains) similar to iGluRs. Tm/DmIR93a contains only the PBP2_iGluR domain similar to other IRs.

**Figure 4.**
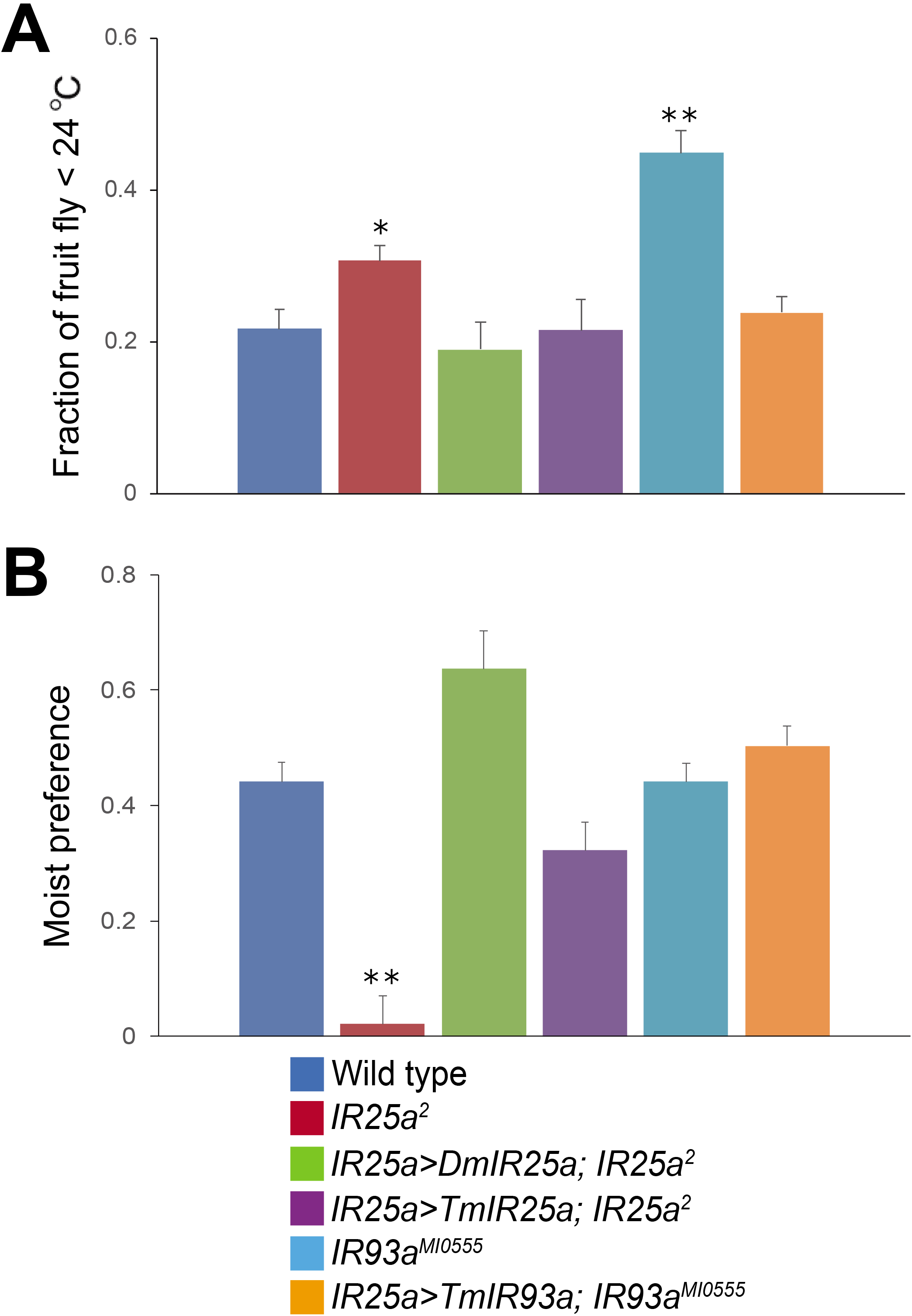
*TmIR25a* and *TmIR93a* rescue the behavioral defects of *Drosophila melanogaster IR25a* and *IR93a* mutants. (A) The fraction of wild type, *IR25a^2^, IR25a^2^* expressing either *DmIR25a (IR25a>DmIR25a; IR25a^2^*) or *TmIR25a (IR25a>TmIR25a; IR25a^2^*), *IR93a^MI0555^*, and *IR93a^MI0555^* expressing *TmIR93a (IR25a>TmIR93a; IR93a^MI0555^*) under *IR25a-Gal4* in the area < 24 °C of the thermal gradient. The recording was repeated 7-21 times for each genotype. The mean value with error bar (± SEM) is shown for each genotype. Asterisks (* and **) are significantly different from wild type, and *P*-values for *IR25a^2^* and *IR93d^M10555^* are < 0.03 and < 0.000002, respectively. (B) Moist preference (70 over 20 % humidity) of fruit flies of above genotypes is shown. The recording was repeated 3-9 times for each genotype. The mean value with error bar (± SEM) is shown for each genotype. Asterisks (**) is significantly different from wild type (*P*-value < 0.000002).

We then tested the humidity preferences of the fruit fly stocks described above. Wild type flies preferred high (saturated NaCl, 70%) over low (saturated LiCl, 20%) humidity but this preference was significantly impaired in *IR25a* and *IR93a* mutants, as previously reported ^33,35,36,38^. However, we did not detect humidity preference defect with *IR93a* mutant. Expression of *DmIR25a* or *TmIR25a* was able to rescue the humidity preference defect of the mutant fly (Fig. 4B). These results demonstrate that the structures and physiological functions of IR25a and IR93a are highly conserved between *D. melanogaster* and *T. mercedesae*.

## Discussion

### Morphology and structure of the *T. mercedesae* sensory organ

We aimed to identify a sensilla-rich sensory organ in the body of *T. mercedesae* using SEM and found two such organs, one on the mouth parts and the other on the dorsal side of the foreleg tarsus. The latter is comparable to Haller’s organ in ticks, which is considered to be responsible for detecting humidity, temperature, and odor ^39,40^. Similar sensory organs have also been identified in the foreleg tarsi of the mites *Dermanyssus prognephilus* ^41^, *Dermanyssus gallinae* ^42^, and *V. destructor* ^26,27^. Thus, acarids are likely to share the same mechanisms for sensory perception. Nevertheless, structural diversity exists between different species. For example, *V. destructor* has nine large sensilla (R1-9) at the periphery and nine small sensilla (S1-9) on the inside of the sensory organ *^26,27^*. The sensory organ of *T. mercedesae* did not have such organization and the localization of small and long sensilla was also random (Fig. 1). The existence of four different types of sensilla appears to be shared between *T. mercedesae* and *D. gallinae*, suggesting that the mite sensory organ could respond to mechanical stimuli, humidity, temperature, and odor. Electrophysiological characterization of each sensillum is, of course, necessary to support this hypothesis.

### *T. mercedesae* sensory organ enriched with mRNAs for sensory proteins and proteins necessary for ciliary biogenesis/transport

We sought to identify mRNAs differentially expressed in the forelegs of mites as candidates for those expressed in the sensory organs. Although we have no direct evidence to show that these mRNAs are indeed expressed in the sensory organ, their specific existence in the forelegs, as well as the identified DEGs, support this approach. The same method was used with two tick species, *Dermacentor variabilis* and *Ixodes scapularis*, to identify mRNAs associated with the Haller’s organ ^43,44^ Our results to show the enrichment of *TmIR25a* and *TmIR93a* mRNAs in the forelegs of *T. mercedesae* are consistent with the results for *I. scapularis* ^44^ Eliash et al. ^45^ also reported that the *V. destructor* homolog of IR25a (this may not be the ortholog since it does not have the N-terminal LIVBP domain) was highly expressed in the forelegs. These results suggest that IR25a and IR93a may represent the major thermo- and hygroreceptors in acarids, based on their physiological roles in fruit flies. This hypothesis was further supported by our finding that TmIR25a rescued the defective thermo- and hygrosensations in *D. melanogaster IR25a* mutant and TmIR93a rescued the defective thermosensation in *D. melanogaster IR93a* mutant (Fig. 4). It is notable that not only the structure, but also the physiological roles, have been deeply conserved during Arthropod evolution ^30^. *T. mercedesae* also showed high expression of two acarid-specific IR mRNAs, Tm15230 and Tm15243, in the forelegs and these may function as chemoreceptors. Two (Tm03548 and Tm05586) and eight GR mRNAs were highly expressed in the sensory organs of *T. mercedesae* and *I. scapularis*, respectively ^44^ However, these GRs do not appear to be orthologs and Josek et al. ^44^ reported that the expression of other *I. scapularis* GRs was too low to make a comparison between the forelegs and hindlegs. Furthermore, most of the IRs and GRs have expanded in Acari in a lineage-specific manner ^44,46^. Except for Gr28b in *D. melanogaster*, which has an important role in thermosensation ^47^, GRs are generally considered to function as chemoreceptors. Thus, the above two IRs and two GRs (four in total) of *T. mercedesae* may detect, for example, a few odorants/tastants derived from honey bee adults, larva, and larval food. This is consistent with the finding that the numbers of IR and GR genes in parasitic *T. mercedesae* were dramatically reduced compared to those in “free-living” mites/ticks ^37^ The TRPA1 channel was also enriched in the forelegs and may function as a sensor to detect nociceptive stimuli (temperature and chemicals) for avoidance, as previously reported ^48–50^. In summary, *T. mercedesae* may depend on IR25a, IR93a, and TRPA1 for thermosensation, IR25a and IR93a for hygrosensation, and two acarid-specific IRs (Tm15230 and Tm15243), two acarid-specific GRs (Tm03548 and Tm05586), and TRPA1 for chemosensation.

Another group of proteins enriched in the forelegs is associated with cilium assembly and intraciliary transport processes and includes kinesin, dynein, and intraflagellar transport proteins. Cilia are organelles present on the cell surface that concentrate signaling molecules to organize sensory, developmental, and homeostatic function. Movement of the signaling receptor from the basal body into the cilia requires IFT-A and its exit depends on IFT-B and BBSome ^51^. Many sensilla are present in the sensory organ of *T. mercedesae* (Fig. 1) and sensory neurons associated with the sensilla have a ciliated dendrite, which requires the protein complexes described above to control traffic of, for example, sensory proteins. GPCRs are considered to be the major target for intraciliary transport ^52^; however, the four IRs of *T. mercedesae* may also depend on IFT-A, IFT-B, BBSome, and other proteins for transport. Consistent with the presence of few sensilla in the Haller’s organs of two tick species, enrichment of these mRNAs was not observed ^39^ In contrast to Carr et al. ^39^, we did not observe high expression of mRNAs for the downstream signaling pathway components of sensory proteins in the forelegs of *T. mercedesae*.

Our study uncovers the ancient roles of IR25a and IR93a in thermo- and hygrosensation of arthropods. We also found the potential roles of evolutionarily conserved intraciliary transport proteins for the entry and exit of sensory proteins in the ciliated dendrites of sensory neurons. The functional disruption of these proteins could be considered as an effective method to control honey bee parasitic mites as well as other mites/ticks that represent major pests for plants and animals.

## Materials and Methods

### Mite sampling

*T. mercedesae* infested honey bee colonies were obtained from a local beekeeper in Suzhou, China. Adult females of *T. mercedesae* were collected from the capped brood cells and dissected under a light microscope using fine forceps. The collected mites were directly used for all experiments and kept together with honey bee pupae in 33 °C incubator when necessary.

### SEM

A cold field emission gun SEM (Hitachi S-4700, Hitachi Company) was used for characterizing sensory organs of *T. mercedesae*. The whole mites and dissected legs were sprayed with gold alloy first, and then mounted on a conductive adhesive tape. During the observation, each sensillum was assigned with a number to classify the types of sensilla.

### RNA-seq

Total RNA was extracted from the forelegs, hindlegs, and main bodies of 50 adult females of *T. mercedesae* using TRI Reagent (Sigma). High-quality RNA samples in duplicate were then sequenced at BGI (Shenzhen, China) using Illumina HiSeq 4000 platform. After sequencing, the raw data were filtered to remove the adaptor sequences, contamination, and low-quality reads by BGI. The Quality control (QC) was further analyzed using FastQC.

### Bioinformatics

The reference genome and annotated genes of *T. mercedesae* were first acquired from NCBI (https://www.ncbi.nlm.nih.gov/genome/53919?genome_assembly_id=313451), and then used for building the index by Hisat2–build indexer ^53^. The generated index files were used to align the clean reads of six RNA-seq samples to the reference genome. Subsequently, SAM file outputs from the previous step were sorted using SAMtools ^54^ HTSeq-count ^55^ was further applied to obtain the raw read counts for downstream analysis of identifying the DEGs in *R* (V3.4.3) based Bioconductor edgeR package (V3.20.9) ^56^. DEGs were cut-off by a False Discovery Rate (FDR) at 0.05, and then they were subjected to gene ontology (GO) term enrichment analysis using Blast2GO ^57^. The results of GO enrichment analysis between the forelegs, hindlegs as well as main bodies were cut-off by FDR at 0.05.

### *TmIR25a* and *TmIR93a* cDNA cloning

For *TmIR25a* and *TmIR93a*, the full length cDNAs were obtained by identifying the 5’ and 3’ ends with RACE method. To amplify 5’ end sequence of *TmIR25a*, the following two primers: 5’-GAGTGTTTGTCCAAGTACATTCTCGA-3’ (1st PCR) and 5’-AGTGTTATCACAAGGAGATATGAGATC-3’ (2nd PCR) were used for 5’RACE with SMART RACE kit (TAKARA). The 3’end sequence was determined by 3’RACE using the following two primers: 5’-CCATCAAGAACATCGGTGGTG-3’ (1st PCR) and 5’-GGCCTGCATCACATTAGTGTTC-3’ (2nd PCR). 5’RACE for *TmIR93a* was conducted with two primers, 5’-ATCGAGTGCGATCACAAGCAG-3’ (1st PCR) and 5’-ACTCTCAGATTCCGGATTCACC-3’ (2nd PCR) using 5’-Full RACE Kit (TAKARA). For the 3’ RACE, two following two primers: 5’-GGGCAAACAGGTTACAGCTTC-3’ (1st PCR) and 5’-CCCCAACAGGACCGATCTTAT-3’ (2nd PCR) were used. *TmIR25a* full length cDNA was amplified by nested PCR using the following primer sets: Forward-5’-GCGTGAACACATCAGGCCGCT-3’ and Reverse-5’-CCCACTCGGAACTTCGTGTCG-3’ (1st PCR), Forward-5’-TTT<u>GCGGCCGC</u>T**ATG**TGGGTCCCTTTACGGATCTC-3’ and Reverse-5’-TTT<u>TCTAGA</u>CTGTATCGCCTGGCGGGGTAGTT-3’ (2nd PCR). Similarly, *TmIR93a* full length cDNA was obtained using the following primer sets: Forward-5’-GGGAGAAAGCCGAGCTGGTAA-3’ and Reverse-5’-TTGTGAATGTCGCCGGTATCC-3’ (1st PCR), Forward-5’-TTT<u>GCGGCCGC</u>GAC**ATG**TGGCCTCGACTCATATTT-3’ and Reverse-5’-TTT<u>TCTAGA</u>CTGTATCGCCTGGCGGGGTAGTT-3’ (2nd PCR). The PCR products were digested by NotI and XbaI and cloned into pAc5.1/V5-His vector (Thermo Fisher Scientific) in which *Drosophila melanogaster Act5C* promoter was replaced by CMV promoter for expression of the V5-epitope tagged proteins in HEK293 cells. To generate *UAS-TmIR25a* and *UAS-TmIR93a* transgenic fruit flies, the untagged versions of above expression constructs were first prepared. The EcoRI-XbaI fragment of *TmIR25a* in above construct was replaced with the restriction enzyme digested PCR product obtained with two primers, 5’-GCCACGATGACCAACTGTGAT-3’ and 5’-CGGGCCC<u>TCTAGA</u>**CTA**TTTCTT-3’. The HindIII-XbaI fragment of *TmIR93a* was replaced with the restriction enzyme digested PCR product obtained with two primers, 5’-GGCCAAGCGGTCATCGAGATA-3 ‘ and 5’-GCCC<u>TCTAGA</u>**CTA**GTATCGCCT-3’. The untagged cDNAs were then cloned in pUASTattB ^58^ digested with NotI and XbaI. The accession numbers for *TmIR25a* and *TmIR93a* are LC438511 and LC438512, respectively.

### Western blot

HEK293 cells in 12-well plate were transfected with 1 μg of above expression construct (the V5-epitope tagged version) using 2 μL of Lipofectamine 2000 under OPTI-MEM medium (Thermo Fisher Scientific) for 24 h. The transfected cells were washed once with PBS, and then lysed with 200 μL of SDS-PAGE sample buffer. The cell lysates were sonicated and heated at 60 °C for 5 min. The proteins were separated by 8% SDS-PAGE and transferred to a NC membrane (PALL, 66485) by Pierce 2 Fast Blotter (Thermo Fisher Scientific, B103602038). The membrane was first blocked with 5% BSA/TBST (10 mM Tris-HCl, 150 mM NaCl, and 0.1% Tween 20) for 40 min, and then incubated with rabbit anti-V5-epitope antibody (SIGMA) (1:1000) for 2 h at room temperature. The membrane was washed with TBST for five times (5 min each), and then incubated with IRDye800-conjugated secondary antibody (LI-COR) (1:10,000) for 1.5 h at room temperature under dark and washed as above. The fluorescent signal was detected by Odyssey (LI-COR).

### Fruit fly genetics

*UAS-TmIR25a* and *UAS-TmIR93a* prepared above were integrated at attP2 site on 3rd chromosome. Stocks of *IR25a-GAL4* (BDSC 41728), *IR25a^2^* (BDSC 41737), *UAS-IR25a* (BDSC 41747) and *IR93a^M105555^* (BDSC 42090) were obtained from Bloomington Drosophila Stock Center (BDSC). Using above stocks, we generated *IR25a^2^ UAS-IR25a, IR25a^2^ IR25a-GAL4, IR25a^2^; UAS-TmIR25a, IR25a-GAL4; IR93a^MI05555^*, and *IR93a^MI05555^ UAS-TmIR93a* stocks. The appropriate crosses were made between *Gal4* and *UAS* stocks to test whether *TmIR25a* or *TmIR93a* can rescue the behavioral defects of *IR25a* or *IR93a* mutant.

### Thermotaxis test

To assay the temperature preference of fruit flies, a temperature gradient of 10-40 °C with a slope of 1.07 °C/cm was produced in an aluminum block (27 long × 15 wide × 2.5 cm high) as previously reported ^59^ The temperature gradient was established using a cold circulating water chamber and a hot probe at each end. The aluminum block was covered with moist paper to maintain a uniform relative humidity along the gradient. This paper was divided into 20 observation fields with a black pencil for recording the distribution of fruit flies. A glass plate with three separate lanes was placed 5 mm above the block, creating suitable corridors for fruit flies to migrate. Approximately 30 adult flies (4-5 days old) per lane were placed in the middle of testing arena around 25 °C between the aluminum block and the glass plate, allowed to migrate for 3 h, and photographed every 10 min with a digital camera. When the positions of fruit flies in the apparatus were stabilized between 1.5 to 2.5 h (This time period did not differ between the experimental groups), the number of fruit flies located at the area < 26 °C was counted. Preference index was calculated by the number of flies at < 26 °C/the total number of flies. The preference indexes of all tested groups were statistically analyzed by one-way ANOVA with multiple comparisons followed by Dunnett test. In each test, wild type, mutant, and the recued fruit flies were examined simultaneously. All experiments were performed in a room where the temperature was kept constant at 25°C.

### Humidity preference test

Hygrosensory behavior was assayed as previously reported ^36^. A 12-well cell culture plate was modified to make a well-defined chamber with two spaces. A half of wells was filled with saturated solution of LiCl (20 % humidity at 25 °C) while another half was filled with saturated NaCl (70 % humidity at 25 °C) to maintain stable relative humidity (RH) on the liquid surface in an enclosed space. The plate was then covered by a nylon mesh and closed with a lid matching the plate. In each test, approximately 80 adult flies (4-5 days old) were briefly ice anesthetized and placed at the center of apparatus. The lid was sealed to stabilize RH inside the apparatus. Humidity preference of the fruit flies with different genotypes were recorded using a digital camera and the numbers of flies on each side were recorded manually at 3-5 h after the start of recording. Humidity preference index was calculated by (the number of flies on NaCl side – the number of flies on LiCl side)/total number of flies. The preference indexes of all tested groups were statistically analyzed by one-way ANOVA with multiple comparisons followed by Dunnett test.

---

**Figure S1.**
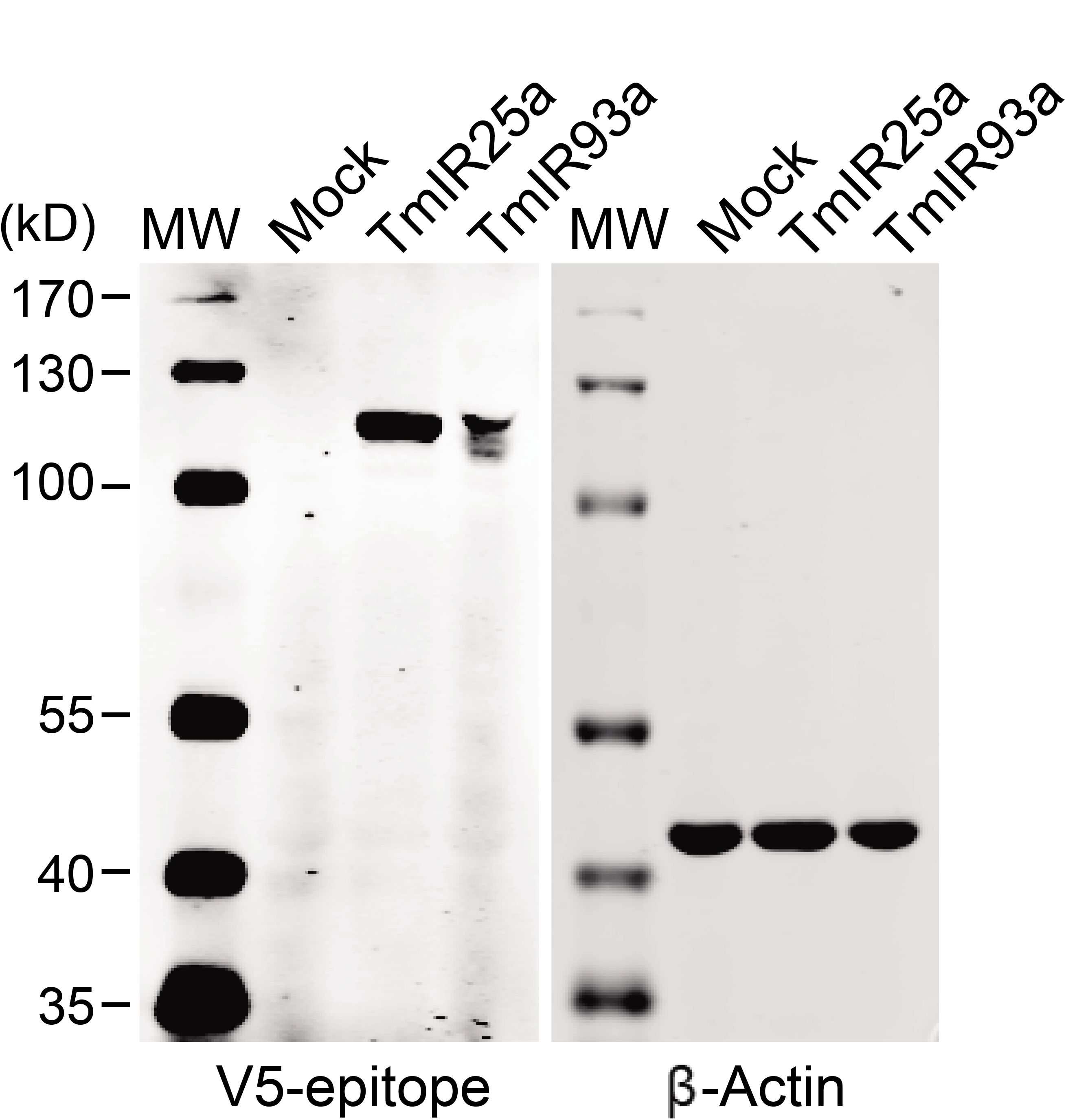
Expression of TmIR25a and TmIR93a proteins. The IR proteins (V5-epitope) and β-actin expressed in HEK293 cells transfected with empty vector (Mock), TmIR25a-, and TmIR93a-expressing constructs were analyzed by western blot. The size (kD) of protein molecular weight marker (MW) is at the left.

