## Supplementary Tables for "Honey bee parasitic mite contains the sensory organ expressing ionotropic receptors with conserved functions": Table S1, S5, S6.docx

**Table S1 Total reads of RNA-seq and the alignment rates to the reference genome.**

| Samples | Total reads | Alignment rate |
| --- | --- | --- |
| Forelegs 1 | 20446207 | 77.4% |
| Forelegs 2 | 20733523 | 52.7% |
| Hindlegs 1 | 20217399 | 83.9% |
| Hindlegs 2 | 20715416 | 46.1% |
| Main bodies 1 | 20379439 | 52.4% |
| Main bodies 2 | 20292712 | 47.8% |

**Table S5 GO terms enriched with genes highly expressed in the forelegs compared with main bodies of *T. mercedesae***

| GO ID | GO name | GO category | FDR | P-value |
| --- | --- | --- | --- | --- |
| GO:0005272 | sodium channel activity | Molecular function | 7.04E-08 | 3.38E-11 |
| GO:0003777 | microtubule motor activity | Molecular function | 3.97E-05 | 1.97E-07 |
| GO:0004970 | ionotropic glutamate receptor activity | Molecular function | 0.00141209 | 1.42E-05 |
| GO:0008017 | microtubule binding | Molecular function | 0.00721176 | 9.81E-05 |
| GO:0004129 | cytochrome-c oxidase activity | Molecular function | 0.00783453 | 1.15E-04 |
| GO:0008121 | ubiquinol-cytochrome-c reductase activity | Molecular function | 0.00950466 | 1.54E-04 |
| GO:0005234 | extracellularly glutamate-gated ion channel activity | Molecular function | 0.03115925 | 5.79E-04 |
| GO:0008417 | fucosyltransferase activity | Molecular function | 0.04673965 | 9.50E-04 |
| GO:0006814 | sodium ion transport | Biological process | 8.38E-06 | 2.68E-08 |
| GO:1905515 | non-motile cilium assembly | Biological process | 0.00287788 | 3.27E-05 |
| GO:0042073 | intraciliary transport | Biological process | 0.00287788 | 3.27E-05 |
| GO:0006120 | mitochondrial electron transport, NADH to ubiquinone | Biological process | 0.00783453 | 1.14E-04 |
| GO:0060401 | cytosolic calcium ion transport | Biological process | 0.02416522 | 4.33E-04 |
| GO:0050907 | detection of chemical stimulus involved in sensory perception | Biological process | 0.02416522 | 4.33E-04 |
| GO:0098662 | inorganic cation transmembrane transport | Biological process | 0.02994775 | 5.51E-04 |
| GO:0006874 | cellular calcium ion homeostasis | Biological process | 0.03931295 | 7.74E-04 |
| GO:0034464 | BBSome | Cellular component | 5.21E-06 | 1.40E-08 |
| GO:0005747 | mitochondrial respiratory chain complex I | Cellular component | 0.00475405 | 6.16E-05 |
| GO:0099512 | supramolecular fiber | Cellular component | 0.0107531 | 1.81E-04 |
| GO:0044430 | cytoskeletal part | Cellular component | 0.02004797 | 3.47E-04 |
| GO:0031514 | motile cilium | Cellular component | 0.02416522 | 4.33E-04 |
| GO:0030992 | intraciliary transport particle B | Cellular component | 0.02416522 | 4.33E-04 |
| GO:0031224 | intrinsic component of membrane | Cellular component | 0.03461975 | 6.59E-04 |
| GO:0015630 | microtubule cytoskeleton | Cellular component | 0.04405208 | 8.81E-04 |

**Table S6 GO terms enriched with genes highly expressed in the hindlegs compared with main bodies of *T. mercedesae***

| GO ID | GO Name | GO Category | FDR | P-Value |
| --- | --- | --- | --- | --- |
| GO:0004129 | cytochrome-c oxidase activity | Molecular function | 1.20E-07 | 1.52E-09 |
| GO:0008137 | NADH dehydrogenase (ubiquinone) activity | Molecular function | 2.11E-06 | 2.95E-08 |
| GO:0046933 | proton-transporting ATP synthase activity, rotational mechanism | Molecular function | 2.86E-06 | 4.08E-08 |
| GO:0008121 | ubiquinol-cytochrome-c reductase activity | Molecular function | 4.07E-05 | 6.98E-07 |
| GO:0016614 | oxidoreductase activity, acting on CH-OH group of donors | Molecular function | 3.14E-04 | 5.85E-06 |
| GO:0048037 | cofactor binding | Molecular function | 9.98E-04 | 2.01E-05 |
| GO:0015986 | ATP synthesis coupled proton transport | Biological process | 2.93E-12 | 1.97E-14 |
| GO:0006099 | tricarboxylic acid cycle | Biological process | 2.79E-09 | 2.81E-11 |
| GO:0006120 | mitochondrial electron transport, NADH to ubiquinone | Biological process | 1.23E-08 | 1.39E-10 |
| GO:0090286 | cytoskeletal anchoring at nuclear membrane | Biological process | 4.07E-05 | 6.98E-07 |
| GO:0044782 | cilium organization | Biological process | 3.73E-04 | 6.99E-06 |
| GO:0006122 | mitochondrial electron transport, ubiquinol to cytochrome c | Biological process | 6.09E-04 | 1.19E-05 |
| GO:0006072 | glycerol-3-phosphate metabolic process | Biological process | 6.09E-04 | 1.19E-05 |
| GO:0005747 | mitochondrial respiratory chain complex I | Cellular component | 3.82E-12 | 2.76E-14 |
| GO:0070069 | cytochrome complex | Cellular component | 1.73E-05 | 2.72E-07 |
| GO:0045261 | proton-transporting ATP synthase complex, catalytic core F(1) | Cellular component | 4.07E-05 | 6.98E-07 |
| GO:0071797 | LUBAC complex | Cellular component | 4.07E-05 | 6.98E-07 |
| GO:0034993 | meiotic nuclear membrane microtubule tethering complex | Cellular component | 2.18E-04 | 3.98E-06 |
| GO:0000276 | mitochondrial proton-transporting ATP synthase complex, coupling factor F(o) | Cellular component | 6.09E-04 | 1.19E-05 |
